# EFHD1 ablation reduces cardiac mitoflash activation and protects cardiomyocytes from ischemia

**DOI:** 10.1101/2021.10.18.464831

**Authors:** David R. Eberhardt, Sandra H. Lee, Xue Yin, Anthony M. Balynas, Emma C. Rekate, Jackie N. Kraiss, Marisa J. Lang, Maureen A. Walsh, Molly E. Streiff, Andrea C. Corbin, Katsuhiko Funai, Frank B. Sachse, Dipayan Chaudhuri

## Abstract

Altered levels of intracellular calcium (Ca^2+^) is a highly prevalent feature in different forms of cardiac injury, producing changes in contractility, arrhythmias, and mitochondrial dysfunction. In cardiac ischemia-reperfusion injury, mitochondrial Ca^2+^ overload leads to pathological production of reactive oxygen species (ROS), activates the permeability transition, and cardiomyocyte death. Here we investigated the cardiac phenotype caused by deletion of *EF-hand domain-containing protein D_1_* (*Efhd1^-/-^*), a Ca^2+^-binding mitochondrial protein whose function is poorly understood. *Efhd1^-/-^* mice are viable and have no adverse cardiac phenotypes. They feature reductions in basal ROS levels and mitoflash events, both important precursors for mitochondrial injury, though cardiac mitochondria have normal susceptibility to Ca^2+^ overload. Notably, we also find that *Efhd1^-/-^* mice and their cardiomyocytes are resistant to hypoxic injury.

## INTRODUCTION

Mitochondrial dysfunction commonly occurs during the progression of heart failure[1]. In healthy hearts, calcium (Ca^2+^) regulates mitochondrial metabolism, morphology, and trafficking [2, 3]. In heart disease, however, excess Ca^2+^ entry into mitochondria causes increased pathological reactive oxygen species (ROS) production, leading to the mitochondrial permeability transition (MPT) [4, 5]. During the MPT, a large channel opens, collapsing the mitochondrial inner membrane potential (ΔΨ), disrupting mitochondrial function, and causing cardiomyocyte death and heart failure.

We sought to investigate novel targets involved in cardiac mitochondrial Ca^2+^ and ROS signaling. In a recent study, the EF-hand domain-containing protein D1 (EFHD1) was identified as a Ca^2+^-sensor for mitoflash activation [6]. Mitoflashes are transient and discreet mitochondrial depolarizations associated with a respiratory and ROS burst [7, 8]. Increased mitoflash activity is known to presage MPT. Thus, proteins that regulate mitoflash production may constitute novel targets or pathways for therapies to prevent cardiac injury.

There has been limited study of EFHD1. It is a 27 kDa protein consisting of two Ca^2+^-binding EF-hand domains and a protein-binding coiled-coiled domain [9]. In its initial characterization, it was found to localize predominantly to mitochondria, possibly the inner mitochondrial membrane [10], while structurally-similar homologs EFHD2 and allograft inflammatory factor-1 (AIF), localize to the cytoplasm[11]. In humans, variation in EFHD1 predicts circulating liver enzyme levels [12]. EFHD1 is also a marker of differentiated state lost in certain cancers [13–15], and its expression increases when an important component of the mitochondrial anti-oxidant system, superoxide dismutase 2, is downregulated [16]. In studies of *Efhd1^-/-^* mice, reduced basal respiration and ATP production were noted in the dorsal root ganglion [17]. A slight increase in peripheral neuronal cell death was noted, though functional consequences appeared absent. Mitochondrial function was also altered in pro-B immune cells after EFHD1 inhibition, with a shift towards glycolysis [18]. These studies provide a partial picture, which suggests that EFHD1 is involved in mitochondrial metabolism at the interface between ROS and Ca^2+^.

In the present study, we describe the cardiac phenotype of *Efhd1^-/-^* mice. We found that *Efhd1^-/-^* mice displayed no overt cardiac pathology, while ROS and mitoflash activity was reduced. We also found that *Efhd1^-/-^* mice are resistant to hypoxia and cell death due to ischemia at baseline. The paucity of baseline phenotypes present in *Efhd1^-/-^* animals suggests it may be safe to target this molecule or its regulatory pathway for cardioprotection.

## METHODS

### Animal handling and genotyping

All animal procedures have been reviewed and approved by the Institutional Animal Care and Use Committee at the University of Utah. *Efhd1^-/-^* mice were obtained from the Jackson Laboratory (Bar Harbor, ME, C57BL/6NJ-Efhd1^*em1(IMPC)J*^/Mmjax). Animals were kept on a C57BL/6NJ background. Animals were housed under standard conditions and allowed free access to food and water. Mice were euthanized via CO_2_ inhalation, with an intermediate check at 4 mins.

### Echocardiography

Mice were initially sedated with 2% inhaled isoflurane (Vet One, NDC13985-046-60) anesthetic. They were restrained in the supine position and fur was cleared from the chest with a depilatory gel, after which anesthesia was reduced. Warmed ultrasound gel was applied to the animal and 2D, M-mode and Doppler images were recorded in short-axis at the level of the papillary muscles, using a Vevo 2100 ultrasound machine equipped with a 55-MHz probe (Visual Sonics, Toronto, Ontario, Canada).

### Mitochondrial isolation

Mice were euthanized as described above and the organs were removed and washed in ice-cold phosphate buffered saline (PBS). Hearts were rapidly dissected to remove atria and great vessels, and then placed in ice-cold initial medium containing (mM): 225 mannitol, 70 sucrose, 5 HEPES, 1 EGTA, and 0.1% (w/v) bovine serum albumin (BSA), and 1 μM thapsigargin (TG) (pH to 7.2 with KOH; osmolality to 290-310 mOsm/L). Tissue was homogenized using a Potter-Elvehjem tissue grinder attached to an overhead stirrer (IKA, Wilmington, NC) for 10-15 strokes at 180 rpm. The homogenate was centrifuged at 700 X *g* for 7 minutes and the tissue pellet was discarded. The supernatant was then centrifuged at 8500 X *g* for 10 minutes to obtain a mitochondrial fraction. The mitochondrial pellet was washed twice with and carefully resuspended using a pipette tip where the end had been removed in imaging solution containing (mM): 125 KCL, 20 HEPES, 5 K_2_HPO_4_, 1 MgCl_2_, and 10 μM EGTA (pH to 7.2 with KOH, osmolality 290-300 mOsm/L). Protein concentration was quantified using the Pierce 660 nm Protein Assay Reagent (Thermo Fisher, Waltham, MA) according to the manufacturer’s directions. In each subsequent assay, we obtained parallel data from knockout and littermate or age- and sex-matched C57BL/6NJ WT controls processed together.

### Isolation of other organelles

For differential centrifugation experiments to study non-mitochondrial organelles, we followed a secondary protocol described previously [19]. Mouse organs were harvested and homogenized as described above for mitochondrial isolation. The homogenate was first spun at 600 x g for 3 minutes at 4°C. The pellet contained nuclei and cell debris. The supernatant was taken and spun at 6,000 x g for 10 minutes. The pellet contained the mitochondrial and lysosomal fraction. The supernatant was taken and spun at 40,000 x g for 30 minutes using a Beckman Coulter® Optima™ MAX-XP bench-top Ultracentrifuge. The pellet contained microsomes from the ER. The supernatant was taken and spun at 100,000 x g for 90 minutes. The pellet contained light microsomes from the endoplasmic reticulum (ER) and ribosomes. The supernatant contained the cytosolic fraction. Purity of each fraction was confirmed via Western Blot using antibodies against known organelle marker proteins.

### Isolation of mitochondrial-associated membranes (MAMs)

MAMs were isolated as described previously [20]. Briefly, the crude mitochondrial pellet was obtained. The crude mitochondria were then layered on top of Percoll medium, consisting of (mM): 225 mannitol, 25 HEPES, 1 EGTA, 30% Percoll (pH 7.4). The crude mitochondria and Percoll medium were spun at 95,000 x g for 30 minutes at 4°C. After the spin, a dense band containing pure mitochondria was located at the bottom of the tube, and a dense band containing MAM was visible above the mitochondrial band. The bands were extracted and spun at 6,300 x g for 10 minutes at 4°C. The mitochondrial fraction was then collected. The MAM fraction was spun again at 100,000 x g for 1 hour and the pure MAM collected. Purity of each fraction was confirmed via Western Blot using antibodies against known organelle marker proteins.

### Proteinase K assay

Mitochondrial fractions were isolated as described above. Mitochondria were resuspended in a sucrose buffer (200 mM sucrose, 10 mM Tris-MOPS, 1 mM EGTA-Tris) to a final concentration of 1 mg/ml. 20 μg of mitochondria were treated with digitonin (0.0001–1%, Sigma) or RIPA (150 mM NaCl, 50 mM Tris-HCL, 1% (w/v) Triton X-100, 0.5 % (w/v) deoxycholic acid, 0.1 % (w/v) sodium dodecyl sulfate (SDS), pH 7.8) and 100 μg/ml of proteinase K (New England Biolabs, Ipswich, MA) for 15 min at room temperature, in a 30 μl final volume. Proteinase K was then inhibited by addition of 5 mM phenylmethylsulfonyl fluoride. 5 μg of protein was loaded to Tris-glycine gels for Western blotting.

### Western blots

Tissue was lysed in RIPA supplemented with protease and phosphatase inhibitors (Thermo Fisher). Protein concentration was quantified by BCA assay (Thermo Fisher). 5-20 μg of protein from total heart lysates were loaded on polyacrylamide gels and processed as described previously on PVDF membranes[21]. We used the following antibodies: β-actin (ab8224, Abcam, Cambridge, MA, 1:1000 dilution), Calreticulin (D3E6, Cell Signaling Technology [CST], Danvers, MA, 1:1000 dilution), GAPDH (14C10, CST, 1:1000 dilution), MCU (D2Z3B, CST, 1:1000 dilution), NDUFS3 (ac14711, Abcam, 1:1000 dilution), PPIF (ac110324, Abcam, 1:1000 dilution), PRELID1 (ab196275, Abcam, 1:1000 dilution), TOM20 (D8T4N, CST, 1:1000 dilution), voltage-dependent anion channel (VDAC, D73D12, CST, 1:1000 dilution). Band intensity was analyzed using ImageJ[22].

The EFHD1 rabbit polyclonal antibody a rabbit was developed by Pacific Immunology, (Ramona, CA). Rabbits were immunized with the EFHD1 peptide sequence: Cys-EQEERKREEEARRLRQAAFRELKAAFSA (Mouse *Efhd1* 212-240). The EFHD1 polyclonal antibody was used at a concentration of 1:500 for Western blot analysis.

### ΔΨ imaging

For measurement of ΔΨ, 100 μg of mitochondria were incubated in 100 μL imaging solution supplemented with (mM): 5 L-glutamic acid, 5 L-malic acid and 20 μM tetramethylrhodamine methyl ester (TMRM, Thermo Fisher) for 10 minutes. We collected baseline signal for 1 minute, then injected 5 μM carbonyl cyanide 4-trifluoromethoxyphenylhydrazone (FCCP), and calculated the maximal change in TMRM signal at excitation and emission wavelengths of 548/574 nm[23]. Imaging was performed in 96-well plates on a Cytation 5 microplate reader.

### Cardiomyocyte isolation and imaging

Mice were euthanized with CO_2_ and their hearts subsequently extracted via thoracotomy. The ascending aorta was cannulated on a 25-gauge blunt needle. Cardiomyocytes were obtained by retrograde perfusion with Collagenase II (1.2 g/L, Worthington Biochemical Corporation, Lakewood, NJ) and Protease XIV (0.1 g/L), and stored in a 0.5 mM Ca^2+^ HEPES-buffered saline solution. Isolated cardiomyocytes were resuspended in warm modified Tyrode solution containing (mM): 126 NaCl, 4.4 KCl, 1 MgCl_2_, 1.1 CaCl2, 24 HEPES, 11 glucose (pH 7.4) and loaded with 20 nM TMRM or 5 μM MitoSox (Thermo Fisher) for 30 minutes at 37°C and then placed in glass-bottomed chamber (MatTek, MA) and mounted on the stage of a Zeiss LSM 510 confocal microscope (Peabody, MA). Excitation and emission were at 548/574 nm. All cells were imaged at the same power and gain settings. Image analysis was performed using Cell Profiler [24].

### Mitoflash imaging

Cardiomyocytes were isolated as above and subsequently placed into KHB buffer containing (mM): 138.2 NaCl, 0.0037, 0.25 CaCl_2_, 0.0012 KH_2_PO_4_, 1.2 MgSO_4_.7 H_2_O, 15 Glucose, 21.85 HEPES (1% BSA, pH 7.2) supplemented with 50 nM TMRM and incubated for 30 minutes at 37°C. Cardiomyocytes were then placed in a glass-bottomed chamber (MatTek, MA) on the stage of a Zeiss LSM 510 confocal microscope (Zeiss, Oberkochen, Germany). Cells were imaged at excitation and emission 548/574 nm for 100 s at a 512 by 32 pixel resolution and scanning frequency of 1000 Hz. Mitoflashes were identified as a transient loss of brightness in TMRM signal. Individual mitoflashes were counted by eye as described previously [25].

### Cardiac histology and immunohistochemistry

Cardiac extraction from animals was as described above for cardiomyocyte isolation. The heart was subsequently incubated in a fixative solution containing 4% paraformaldehyde in PBS for 48 hours at 4°C and then placed in a 70% ethanol solution. The samples were embedded in paraffin, cut, and stained by the Research Histology core at the Huntsman Cancer Institute (University of Utah) and ARUP Research Institute (Salt Lake City, Utah). Staining was for Masson’s trichrome.

### Calcium retention capacity (CRC)

Imaging was performed in 96-well plates on a Cytation 5 microplate reader. 100 μg of mitochondria were incubated in 100 μL of imaging solution supplemented with (mM): 5 L-glutamic acid, 5 L-malic acid, and 20 μM TMRM. Excitation and emission wavelengths were 548/574 nm. Bolus injections of 5 μM CaCl2 were injected into each well every 5 minutes for 60 minutes. CRC was analyzed by calculating the amount of Ca^2+^ added to each well before an MPT event was observed.

### CLAMS chambers

Mice were housed for 72 h in a four-chamber Oxymax system (CLAMS; Columbus Instruments, Columbus, OH) as previously described[26]. O_2_ and CO_2_ content of the exhaust air from each chamber was compared with ambient air O_2_ and CO_2_ content. Food consumption was monitored by electronic scales, water by electronic sipper tubes, and movement by X/Y laser beam interruption. Respiratory exchange ratio (RER) was calculated as VCO_2_/VO_2_. Work was performed by the Metabolic Phenotyping Core, a part of the Health Sciences Cores at the University of Utah. Analysis was conducted using CalR [27].

### Mitochondrial respiration (OXPHOS)

Pellets from the mitochondrial isolation step were suspended in buffer Z (MES potassium salt; 105 mM, KCl 30 mM, KH_2_PO_4_ 10 mM, MgCl_2_ 5 mM, and BSA 0.5 mg/ml), and analyzed by high-resolution respirometry (Oroboros O_2_k Oxygraphs) to determine substrate-dependent respiration. Respiration was measured in response to the following substrate concentrations: 0.5 mM malate, 5 mM pyruvate, 2 mM ADP, 10 mM succinate, and 1.5 μM FCCP, as described previously[28]. Respiration rates were normalized to total protein content. Respiratory acceptor control ratio (RCR) was calculated as State 3/State 4 respiration [29].

### Ischemia/reperfusion protocol for neonatal mouse cardiomyocytes

A glucose-free, serum-free “ischemic” cell media was made using Dulbecco’s modified eagle’s medium (DMEM) without glucose, L-glutamine, phenol red, sodium pyruvate or sodium bicarbonate (Sigma, D5030) Potassium concentration of the media was doubled by adding 0.4 g/L KCl and pH was adjusted to 6.5 to simulate typical ischemic conditions of hyperkalemia and acidosis. The “ischemic” DMEM was gassed with 95% nitrogen, 5% carbon dioxide for 1 hr prior to use to expel any excess oxygen.

Neonatal mouse ventricular myocytes (NMVMs) were isolated from 1-2 day old mice following Pierce™ primary isolation kit (88281, Thermo Fisher Scientific). After 72 hours in standard culture conditions, the NMVM media was replaced with “ischemic” DMEM and plates were placed in a sealed contained at 37 °C with constant gas flow of 95% nitrogen, 5% carbon dioxide to ensure complete depletion of oxygen from culture. After 4 hours in “ischemia,” NMVMs were washed and normal DMEM was replaced. NMVMs were left in cell culture incubator for 24 hours for a period of “reperfusion.”

For Live/Dead analysis, cells were washed twice in PBS then incubated at room temperature in a PBS solution containing 100 μM 4’,6’-diamidino-2-phenylindole (DAPI) as described previously [30]. Cells were then imaged using a Cytation 5 microplate reader on area-scan mode at excitation and emission of 358/461 nm. The fluorescence levels were normalized to a blank well. A well where the cells had been killed using 10% (v/v) Dimethyl sulfoxide (DMSO) was used as a reference for 100% cell mortality. The fluorescent signal from ischemic cells was compared with non-ischemic controls which were treated in the same way minus exposure to ischemic conditions.

### Statistical analysis

Microsoft Excel and OriginPro (OriginLab) were used for data analysis. We rejected the null hypothesis for P values < 0.05. For comparison tests on samples where n < 15, we established used a Mann-Whitney statistical test. For comparisons on samples > 15, we used a two-tailed, unequal variance, Student’s t-test. For the analysis of contingency tables, we performed a Fisher’s exact test using Prism (GraphPad Software).

## RESULTS

### EFHD1 localizes to the mitochondrial outer membrane and intermembrane space

We obtained a custom-made rabbit polyclonal antibody targeting an internal sequence common to human and mouse EFHD1. In agreement with the initial report [10], we detected EFHD1 at high levels in brain and kidney mitochondria. It was also present at detectable levels in adult (>8 weeks old) heart and skeletal muscle, with somewhat lower levels in lung and liver mitochondria (Figure 1A). We confirmed EFHD1 is enriched in mitochondrial fractions following differential centrifugation. Notably, we also found EFHD1 in both the ER and cytosolic fractions (Figure 1B).

**Figure 1.**
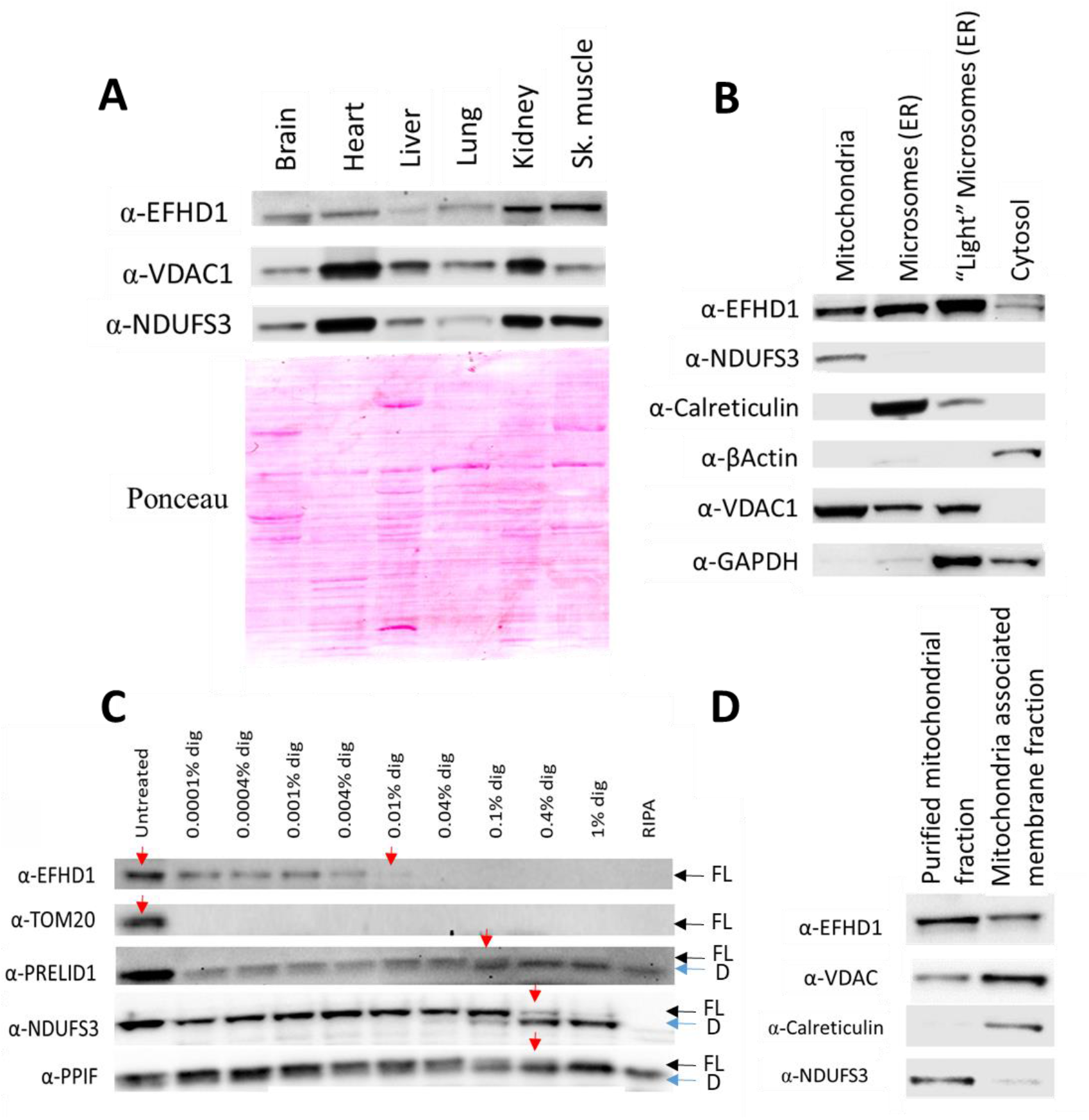
Tissue and subcellular distribution of EFHD1. (A) Western blot analysis of EFHD1 in isolated mitochondria from various organs. Ponceau stain shown as loading control for total protein. VDAC1 and NDUFS3 show are mitochondrial outer and inner membrane proteins. (B) Western blot analysis of cellular fractions separated via differential centrifugation. Fractions were tested against known cellular biomarkers: NDUFS3 (mitochondrial inner membrane); Calreticulin (ER); β-Actin (cytosol); VDAC1 (outer mitochondrial membrane); and GAPDH (cytoplasm). (C) Western blot analysis of Proteinase-K assay showing the breakdown of EFHD1 in isolated mitochondria following treatment with increasing concentrations of digitonin. Fractions were tested against known cellular biomarkers: TOM20 (outer mitochondrial membrane); PRELID1 (mitochondrial inter-membrane space); NDUFS3 (inner mitochondrial membrane); and PPIF (mitochondrial matrix). For PRELID1, NDUFS3, and PPIF, proteinase K accessibility is revealed by shift from full-length (FL) protein to a lower molecular weight digestion product (D). Red arrows show where proteins were digested. Blots are representative of >3 repeats. (D) Western blot analysis of purified mitochondrial fractions and mitochondria-associated membrane fractions. Fractions were tested against known cellular biomarkers.

Furthermore, and contrary to previous findings, our data suggest that EFHD1 is not a mitochondrial inner membrane protein. To examine subcellular location, we performed a proteinase K assay. In this test, mitochondria isolated from mouse kidney were exposed to increasing concentrations of digitonin and assayed for enzymatic digestion with proteinase K [31, 32]. At low concentrations, digitonin will only permeabilize the outer mitochondrial membrane, exposing proteins at the outer membrane and within the intermembrane space to degradation. At higher concentrations, digitonin will also break down the inner mitochondrial membrane, exposing proteins at the inner membrane and within the matrix to degradation. We found that EFHD1 underwent an initial large degradation similar to the outer mitochondrial membrane protein TOMM20 and then a secondary breakdown mimicking PRELID1, a protein in the intermembrane space (Figure 1C). This suggests that EFHD1 resides on the outer membrane and within the intermembrane space of mitochondria. To further confirm this, we used differential ultracentrifugation [20] to isolate mitochondrial membranes which associate with the ER, which constitute purely outer membrane fractions. We found that EFHD1 was present in these ER-associated mitochondrial membranes in a similar way to VDAC1, an outer mitochondrial membrane protein (Figure 1D). These data all suggest that EFHD1 is present within the cytosol and associates with the mitochondrial outer membrane and intermembrane space.

### *Efhd1^-/-^* mice are viable

Having established the localization of EFHD1, we characterized basic cardiac phenotypes of the whole-body *Efhd1^-/-^* mouse, obtained from the Jackson Laboratory (Figure. 2A). Our antibody effectively distinguished wild-type versus *Efhd1^-/-^* in heart, liver and kidney tissue (Figure 2B-D). We found no difference in physical size between age- and sex-matched WT and *Efhd1^-/-^* mice (Figure 2E). Moreover, we also measured body weight and tibia length of WT and *Efhd1-/-* mice and found no difference in these features (Figures 2F and 2G).

**Figure 2.**
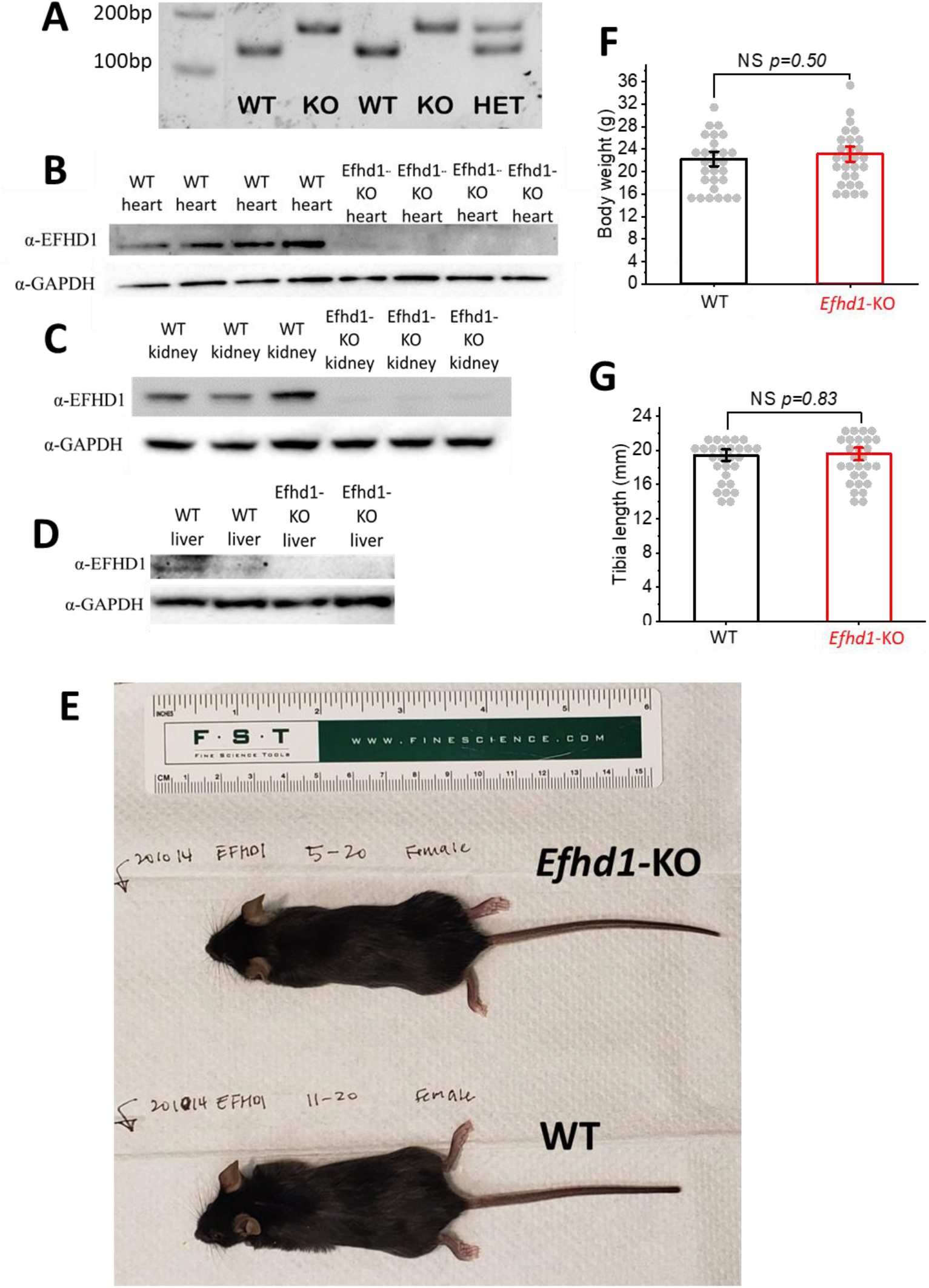
Basic physiology of *Efhd1^-/-^* mice is unaltered. (A) Genotyping data confirming WT, *Efhd1^-/-^* (B-D) Western blot analysis confirming that EFHD1 is absent in heart (A), kidney (B) and liver (C) tissue in *Efhd1^-/-^* mice. (KO) and heterozygous mice. (E) Representative photo showing size similarity between WT (bottom) and *Efhd1^-/-^* (top) mice. (F, G) No difference was observed in body weight (E, n=28) or in tibia length (F, n=28). All mice were adults (>8 weeks). Statistics: Student’s *t*-Test.

EFHD1 has previously been reported to alter mitochondrial metabolism and ROS production, so we tested whether loss of EFHD1 altered metabolism or activity at the whole-organism level. We placed WT and *Efhd1^-/-^* mice in Comprehensive Lab Animal Monitoring System (CLAMS) metabolic cages for 72 hours. We found no differences in respiratory parameters (Figure 3A-P), movement (Figure 3Q and 3R), or food intake (Figure 3S). These findings demonstrate that organismal metabolic homeostasis is unaffected at baseline in *Efhd1^-/-^* mice.

**Figure 3.**
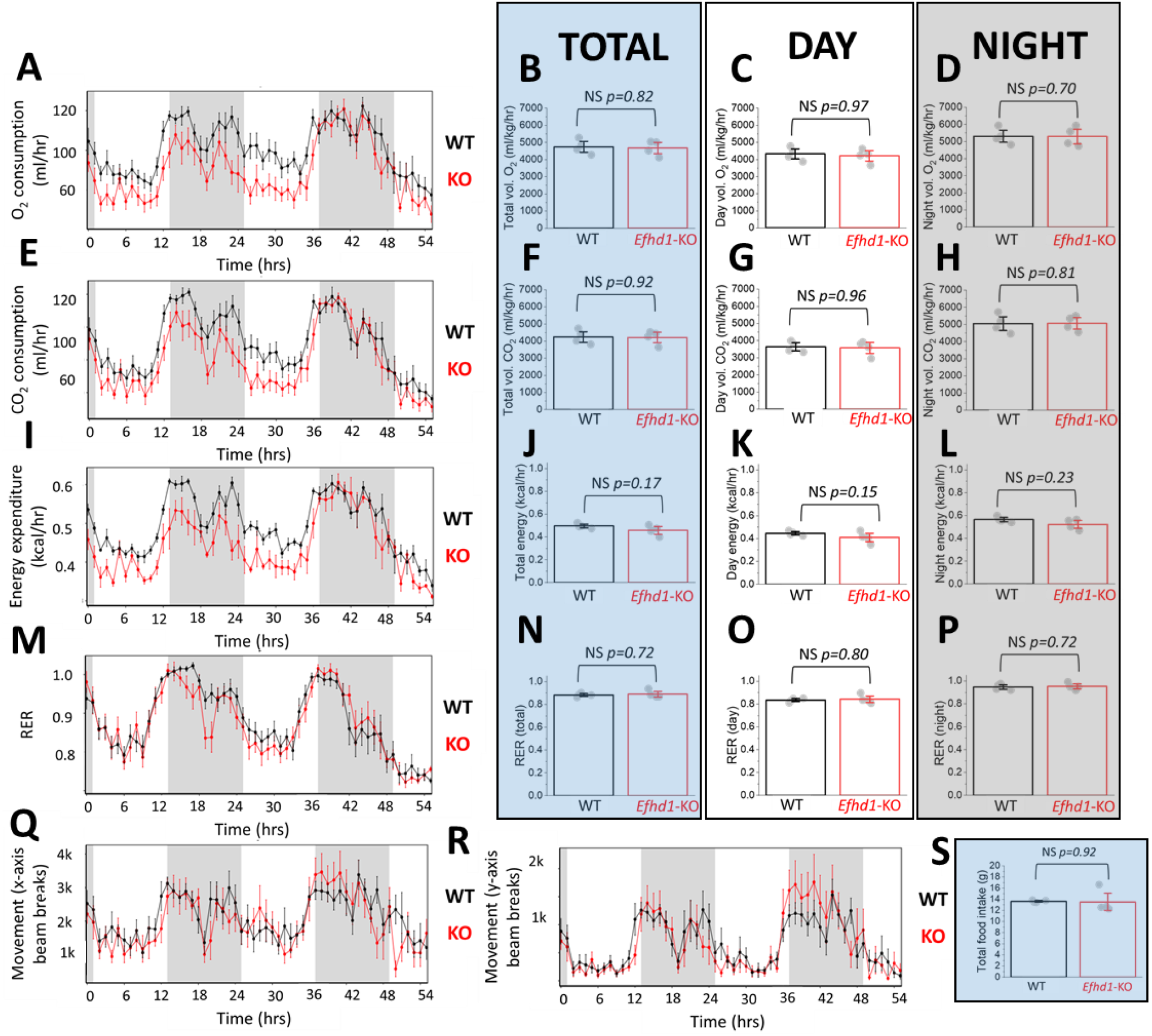
No differences in basal metabolism noted in *Efhd1^-/-^* mice. (A) Line graph showing oxygen consumption over time in mice placed in a CLAMS metabolic chamber for 48 hours. WT mice (n=4) are shown in black. *Efhd1^-/-^* mice (n=4) are shown in red. (B) Comparison of total oxygen consumed in WT (black, n=4) and *Efhd1^-/-^* (red, n=4) mice during 48 hours in a CLAMS chamber. (C, D) Comparison of total oxygen consumed in WT (black, n=4) and *Efhd1^-/-^* (red, n=4) mice during the light/day and dark/night, respectively. (E-H) Same as A-D but for volume of carbon dioxide produced. (I-L) Same as A-D but for energy expended, measured as heat produced. (M-P) Same as A-D but for Respiratory exchange ratio (RCR). (Q, R) Same as A but showing mouse movement measured as the number of times the mouse breaks a beam, locomotor activity (x-axis beam, Q) movement and ambulatory activity (y-axis beam, R), respectively. (S) Comparison of food intake in WT (black, n=4) and *Efhd1^-/-^* (red, n=4) mice placed in a CLAMS metabolic chamber for 48 hours. Throughout the figure, a white background represents data collected during the light/day, a grey background represents data collected during the dark/night, and a blue background represents combined data for day and night. Statistics: Mann Whitney test.

### *Efhd1^-/-^* mice exhibit normal cardiac function

We next focused on cardiac function in the *Efhd1-/-* mice. We performed physical measurements, echocardiography and histology on hearts from WT and *Efhd1-/-* mice. We found no difference in the appearance of hearts from WT and *Efhd1-/-* mice (Figure 4A), and tissue histology was also similar, with no evidence of increased fibrosis (Figure 4B-C). We found no difference in the absolute or normalized weights of the mouse hearts (Figure 4D, F, G). The International Mouse Phenotyping Consortium analyzed electrocardiographic parameters in these mice, and found no difference compared to WT animals [33, 34].

**Figure 4.**
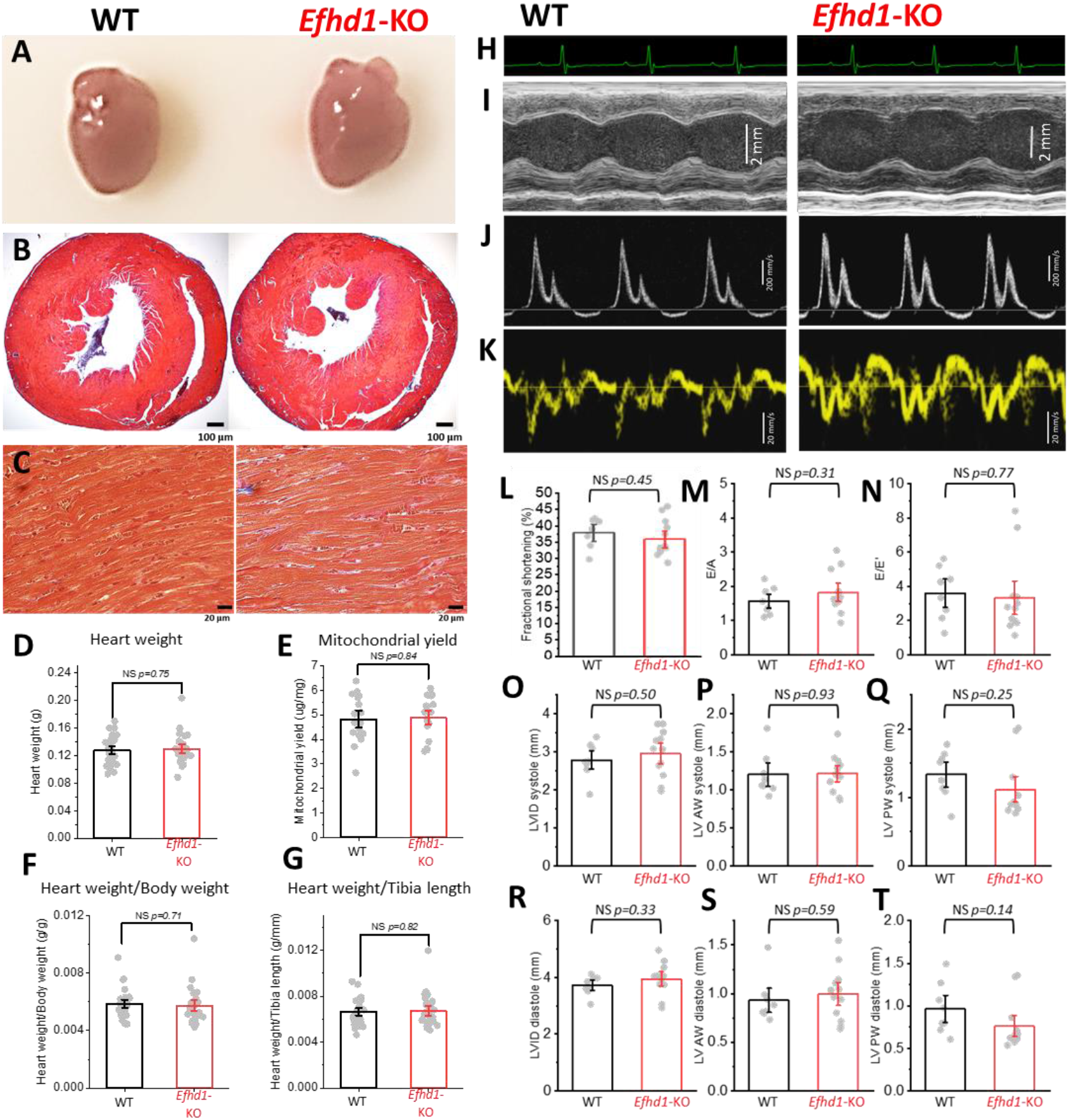
Basic cardiac physiology is unchanged in *Efhd1^-/-^* mice. (A) A representative picture of WT (left) and *Efhd1^-/-^* (right) hearts taken side-by-side showing no difference in size. (B) Histology of transverse slices through WT (left) and *Efhd1^-/-^* (right) hearts. (C) High resolution histology images of WT (left) and *Efhd1^-/-^* (right) hearts treated with Masson’s trichrome staining showing no difference in fibrosis levels (blue staining) in cardiomyocytes of *Efhd1^-/-^* mice compared with WT. Images are representative of 5 WT and 5 *Efhd1^-/-^* mice. (D) Comparison between heart weights of WT (black) and *Efhd1^-/-^* (red) mice (WT: n=28; *Efhd1^-/-^:* n=28). (E) Comparison of mitochondrial yield from WT (black) and *Efhd1^-/-^* (red) mouse hearts (WT: n=17; *Efhd1^-/-^*: n=18). (F) Comparison of Heart weight/Body weight ratio of WT (black) and *Efhd1^-/-^* (red) mice (WT: n=28; *Efhd1^-/-^:* n=28). (G) Comparison of Heart Weight/Tibia length ratio of WT (black) and *Efhd1^-/-^* (red) mice (WT: n=28; *Efhd1^-/-^:* n=28). (H) Representative ECG recordings collected from age-matched WT (left) and *Efhd1^-/-^* (right) mice. (I) Representative m-mode echocardiogram recordings collected from age-matched WT (left) and *Efhd1^-/-^* (right) mice. (J) Representative pulse wave Doppler echocardiogram recordings showing peak velocity blood flow from left ventricular relaxation in early diastole (first peak) and peak velocity flow in late diastole caused by atrial contraction (second peak) collected from age-matched WT (left) and *Efhd1^-/-^* (right) mice. (K) Representative pulse wave Tissue Doppler echocardiogram recordings collected from age-matched WT (left) and *Efhd1^-/-^* (right) mice. H-K were taken from the same WT and *Efhd1^-/-^* mice. (L-T) Comparison of: fractional shortening (L); E/A ratio (M); E/E’ ratio (N); LVID during systole (O); LVAW during systole (P); LVPW during systole (Q); LVID during diastole (R); LVAW during diastole (S); LVPW during diastole (T); in WT (red) and *Efhd1^-/-^* (black) mice measured from echocardiograms. WT: n=8; *Efhd1^-/-^:* n=12. Statistics: Student’s T-Test and Mann-Whitney statistical test.

Next, we tested for differences in basic cardiac function. We performed echocardiograms on male and female age-matched WT and *Efhd1-/-* mice from three separate litters. Representative traces from a WT and *Efhd1-/-* mouse are shown in Figure 4H-K. We found no significant differences in fractional shortening (Figure 4L), E/A ratio (Figure 4M) or in E/E’ ratio (Figure 4N), markers of systolic and diastolic function. We also found no differences between left ventral diameter and wall thickness in both systole (Figure 4O-Q) and diastole (Figure 4R-T). We therefore concluded that the absence of EFHD1 in mice did not result in any overt cardiomyopathy.

We next focused on characterizing the effect of EFHD1 absence on mitochondrial function. We did not see any difference in mitochondrial protein yield between WT and *Efhd1^-/-^* hearts (Figure 4E). Reduced respiration levels have previously been reported in dorsal root ganglion cells from *Efhd1^-/-^* mice [17]. We therefore used an Oroboros O_2_K-FluoRespirometer to measure respiration rates in mitochondria isolated from WT and *Efhd1-/-* mouse hearts. Figure 5A and 5B shows representative traces obtained from WT and *Efhd1-/-* heart mitochondria, respectively. We did not find differences in Complex I-linked respiration (Figure 5C); OXPHOS capacity of complex I, driven by NADH related substrates (Figures 5D and 4E); maximal OXPHOS capacity with convergent input through both complex I and complex II (Figure 5F); maximal convergent respiratory capacity (Figure 5G); or respiratory efficiency, measured as the respiratory control ratio (Figure 5H)[35, 36]. Taken together, this suggests that EFHD1 ablation does not alter basal cardiac mitochondrial function.

**Figure 5.**
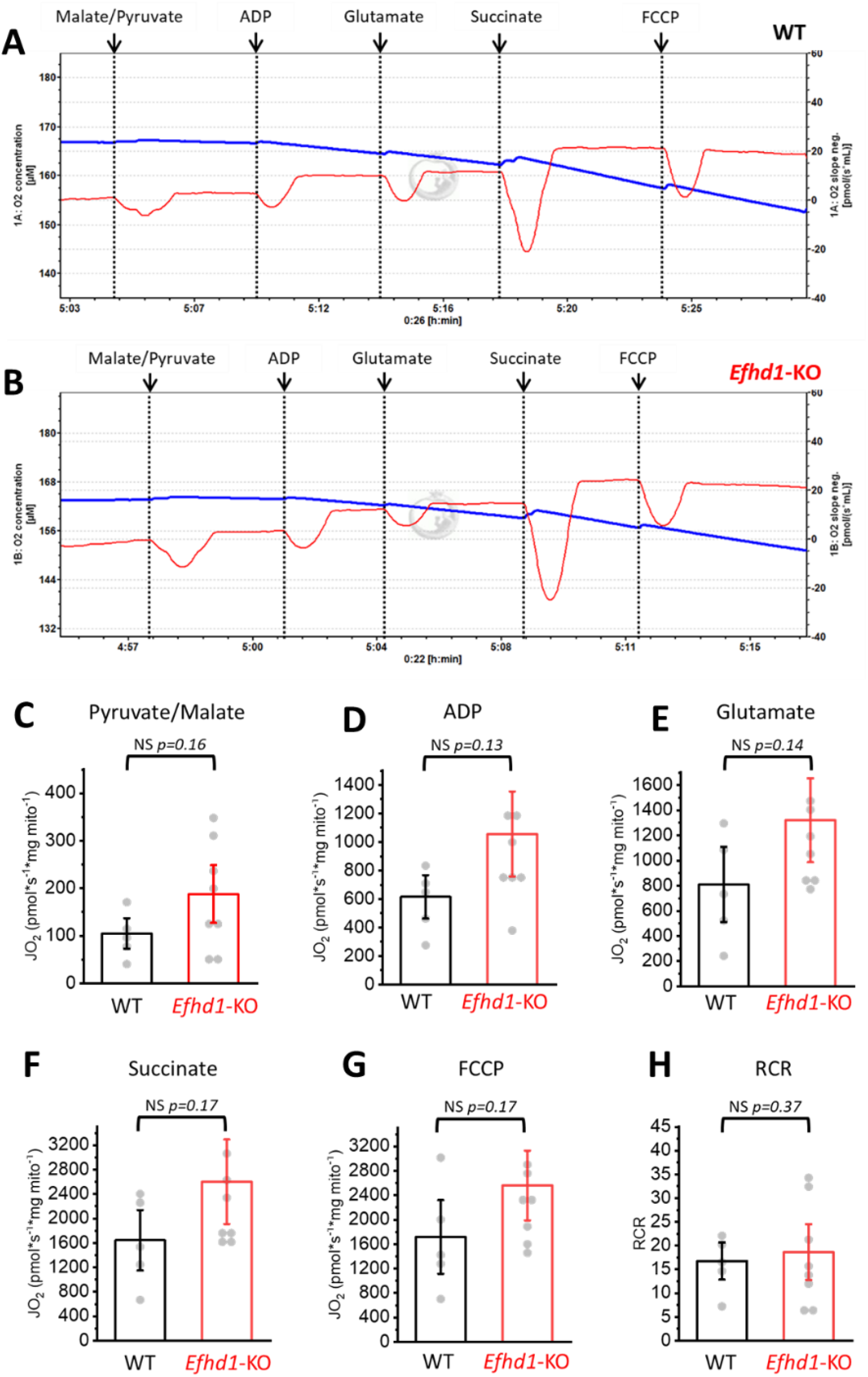
Respiration in mitochondria from *Efhd1^-/-^* mice. (A, B) Representative traces from an Oroboros O_2_k-FluoRespirometer showing oxygen consumption in mitochondria isolated from WT (A) and *Efhd1^-/-^* (B) hearts in response to respiratory substrates. The blue line represents oxygen levels. The red line represents rate of oxygen consumption. (C-G) Comparison of the rate of oxygen consumption of mitochondria from WT (black) and *Efhd1^-/-^* (red) in the presence of respiratory substrates. (H) Comparison of the respiratory acceptor control ratio (RCR) in mitochondria from WT (black) and *Efhd1^-/-^* (red). WT: n=5; *Efhd1^-/-^:* n=8. Statistics: Mann-Whitney statistical test used to compare data in C-H.

### Cardiac *Efhd1-/-* mitochondria are not resistant to Ca^2+^ overload but have reduced ROS and mitoflash levels

Prior experiments in HeLa cells[6] suggested EFHD1 knockdown reduced the frequency of mitoflashes, transient increases in respiration stimulated by increased mitochondrial ROS, which are potential precursors of MPT. Thus, we were interested in testing the effect of deleting EFHD1 on Ca^2+^-overload induced MPT, since EFHD1 is a Ca^2+^ sensor. We used the Ca^2+^retention capacity (CRC) assay, in which purified mitochondria are repeatedly challenged with Ca^2+^ boluses until MPT activation causes depolarization and Ca^2+^ release. We found no difference in cardiac CRC (Figures 6A, B) or expression of mitochondrial Ca^2+^ uptake proteins (Figure 6D, E), between WT and *Efhd1*^-/-^ mitochondria.

**Figure 6.**
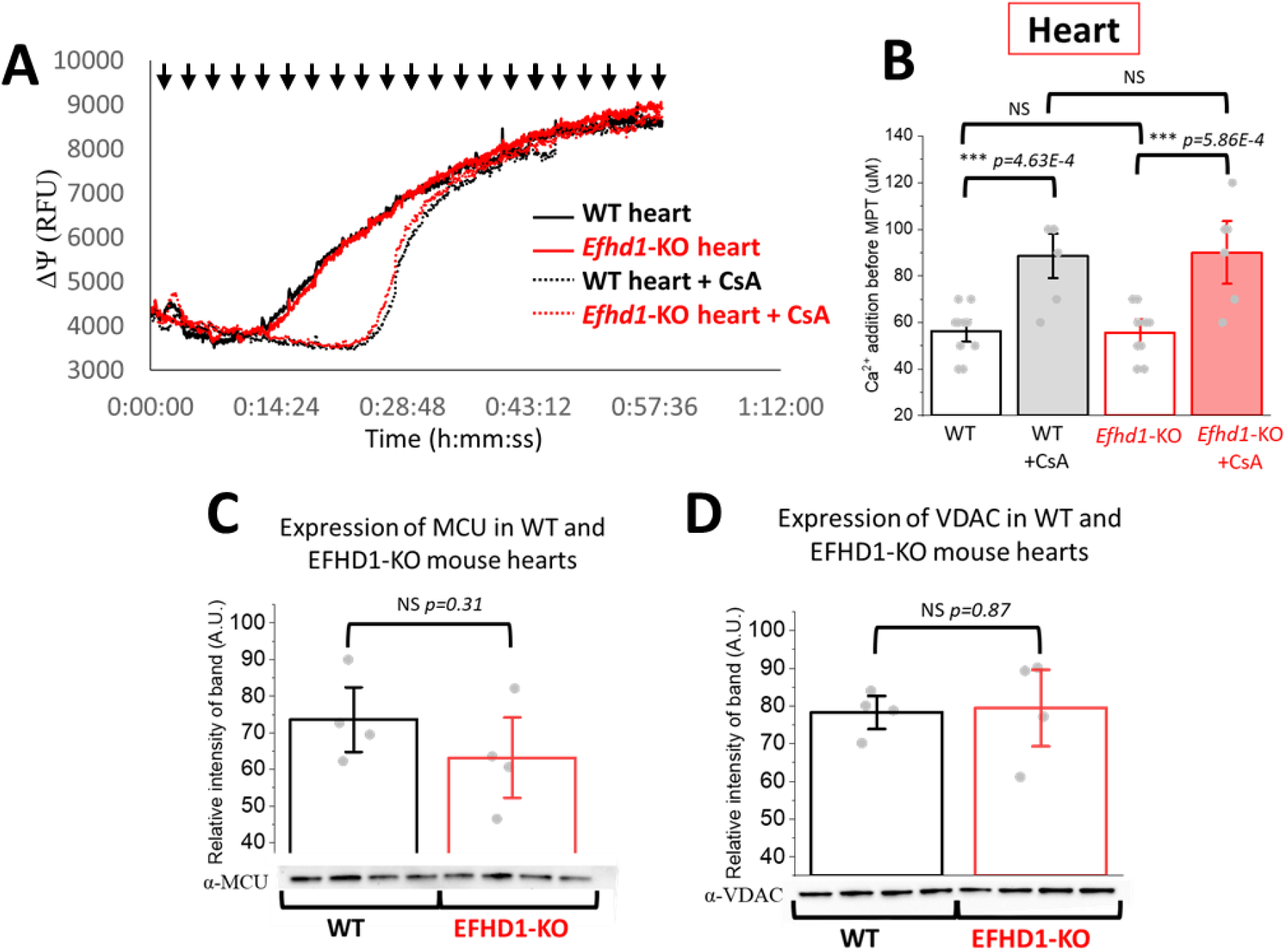
*Efhd1^-/-^* mice have unaltered calcium retention capacity in mouse heart mitochondria. (A) A representative calcium retention trace of mitochondria isolated from WT (black) and *Efhd1^-/-^* (red) mouse hearts in the presence and absence of 10 μM Cyclosporin A (CsA). Black arrows indicate 5 μM Ca^2+^ injections. Mitochondrial viability was measured as ΔΨ/TMRM fluorescence. (B) Comparison of calcium retention before the triggering of an MPT event in WT and *Efhd1^-/-^* mice in the presence and absence of CsA. WT: n=11; WT+CsA: n=7; *Efhd1^-/-^:* n=9; *Efhd1^-/-^* +CsA: n=6. Mitochondrial viability was measured as ΔΨ/TMRM fluorescence. (C) Relative expression of the mitochondrial calcium uniporter (MCU) in WT and *Efhd1^-/-^* mouse hearts. (E) Relative expression of the voltage-dependent anion channel (VDAC) in WT and *Efhd1^-/-^* mouse hearts. Statistics: (B) 1-way ANOVA with a Bonferroni posttest, (C, D) Mann-Whitney statistical test.

Since ROS can trigger MPT and mitochondrial dysfunction independent of Ca^2+^ overload, we next examined these by measuring mitoflashes and ROS levels. Mitoflashes were manually counted over 100s as described previously [25]. In concordance with the prior study, we found that *Efhd1-/-* cardiomyocytes exhibited significantly fewer mitoflashes than WT, with far more cells producing no mitoflashes at all (Figure 7A, B). Moreover, ROS-sensitive MitoSox dye levels were significantly reduced in *Efhd1*^-/-^ cardiac mitochondria (Figure 7E, F) consistent with the reduction in mitoflashes.

**Figure 7.**
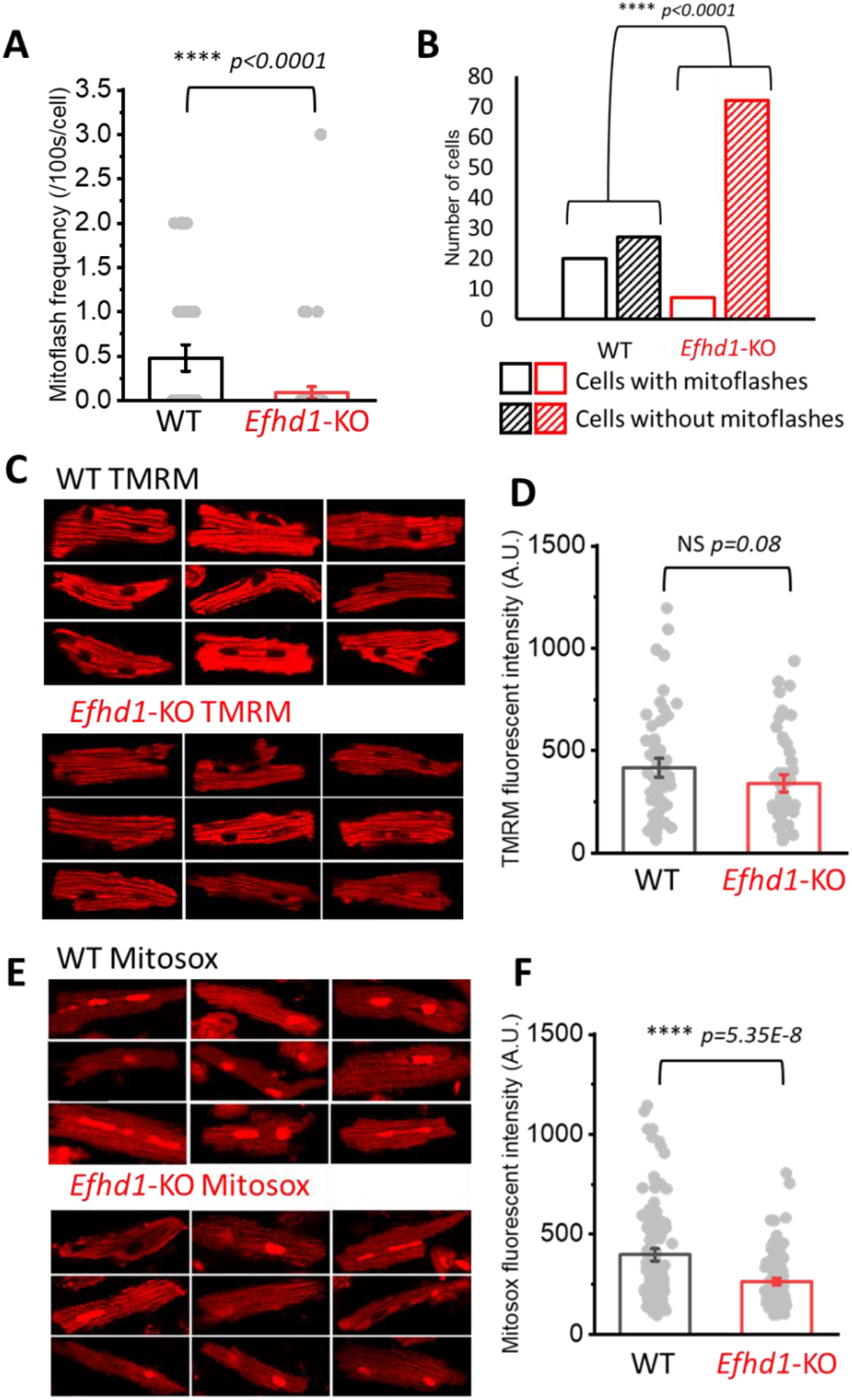
ROS levels and mitoflash frequency is decreased in *Efhd1^-/-^* mouse cardiomyocytes. (A) Comparison of mitoflash frequency between cardiomyocytes from WT (n=50) and *Efhd1^-/-^* (n=79) mice. (B) Comparing the number of WT (black) and *Efhd1^-/-^* (red) cardiomyocytes with mitoflash events against those without. (C) Representative images of cardiomyocytes from WT (top) and *Efhd1^-/-^* (bottom) mice stained with TMRM. (D) Comparison of ΔΨ measured as relative TMRM fluorescence in WT (black, n=69) and *Efhd1^-/-^* (red, n=55). (E) Representative images of cardiomyocytes from WT (top) and *Efhd1^-/-^* (bottom) mice stained with MitoSox. (F) Comparison of ROS levels measured as relative MitoSox fluorescence in WT (black, n=131) and *Efhd1^-/-^* (red, n=125). Statistics: (A, D, F) Student’s T-Test; (B) Fisher Exact test.

### *Efhd1^-/-^* mice are resistant to hypoxia

In performing our assays, we noticed that *Efhd1-/-* mice took much longer to euthanize with CO_2_ when compared with their WT counterparts. Our protocol involves exposing mice to CO_2_ continuously in an enclosed chamber, checking their status at 4 minutes, which in littermate WT mice leads to 100% mortality. However, we were surprised to find that 50% of our *Efhd1^-/-^* mice were still alive at 4 mins (Figure 8A), often requiring 7 to 8 minutes. To examine if a similar effect could be reproduced in cardiac tissue, we isolated neonatal mouse ventricular myocytes (NMVMs) from *Efhd1-/-* heart. We collected NMVMs from 1-day old WT and *Efhd1^-/-^* pups and exposed half of the cells to a deoxygenated, high-potassium, low-pH media for 4 hours, mimicking ischemic conditions, while the other half remained in normal media. Subsequently, cells were grown for 24 hours in regular oxygenated media, to mimic reperfusion, after which the amount of dead cells in each condition was quantified. We found that NMVMs from *Efhd1^-/-^* were twice as resistant to cell death when exposed to ischemic conditions compared to their WT counterparts (Figure 8B). Taken together, our results suggest *Efhd1^-/-^* mice are resistant to cardiomyocyte cell death, possibly through a mechanism involving mitochondrial ROS production.

**Figure 8.**
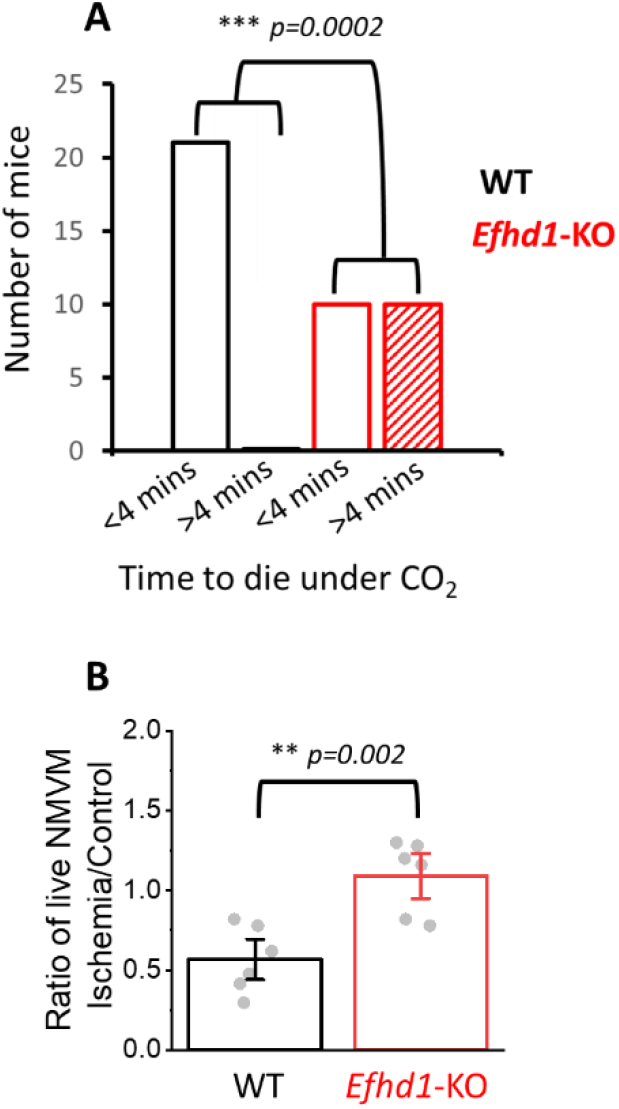
*Efhd_1_^-/^-* mice are resistant to hypoxia. (A) Comparing the number of WT (black) and *Efhd1^-/-^* (red) mice which died in less than 4 mins exposure to CO_2_ with the number which took longer than 4 mins to die (WT n=21; *Efhd1^-/-^* n=20). (B) Comparison of resistance to simulated ischemia-reperfusion in cultured neonatal mouse ventricular myocytes (NMVMs) isolated from 1-day old WT (black) and *Efhd1^-/-^* (red) mouse pups. WT: N=6; *Efhd1^-/-^:* N=6, where each “N” represents NMVMs isolated from one litter (3-10 pups). Statistics: (A) Fisher Exact; (B) Mann-Whitney statistical test.

## DISCUSSION

Here, we present the initial characterization of the baseline cardiac phenotype of *Efhd1*^-/-^ mice. The major conclusions are that *Efhd1^-/-^* mice (1) do no display any obvious organismal or cardiac pathology at baseline, (2) cardiomyocyte ROS and ROS-dependent mitoflashes are reduced, and (3) at the organismal and cardiomyocyte level, they are resistant to hypoxic cell death.

### EFHD1 resides on the outer membrane and intermembrane space of mitochondria

In this study, we used a knockout-validated EFHD1 antibody to demonstrate that that EFHD1 resides primarily on the outer mitochondrial membrane, with lower protein levels being present in the intermembrane space (Figure 1C). This was confirmed using differential centrifugation, a proteinase K protection assay, and purification of ER-associated mitochondrial outer membranes. EFHD1 possesses no transmembrane domains, so it is likely only associated with membranes, rather than an integral component. At first glance, this finding appears contrary to previously findings, which report EFHD1 is an inner membrane protein [10]. However, there has been little investigation of EFHD1 function, and its subcellular localization has been directly examined in only the initial report [10]. In that report, inner membrane localization was assigned based on an immunostaining showing colocalization with a mitochondrial marker, and an immunogold assay showing enrichment in inner membrane on electron microscopy. However, there are two strong caveats related to that evidence. First, even in that report, the authors described substantial EFHD1 staining outside the inner membrane in several other figures. In fluorescent immunostaining, EFHD1 was seen in cytoplasm, neurites, and growth cones. In immunogold staining, although there was greatest labeling of the inner membrane, there is also a diffuse pattern of staining throughout mitochondria and other locations. Second, the authors report that, on western blotting, their rabbit polyclonal antibody detects several non-specific bands. Thus, it is unclear if the staining seen with fluorescence or electron microscopy is entirely specific to EFHD1. Because we also found substantial non-specific bands using our antibody, we limited our analysis to western blots, where we have validated the specific band using *Efhd1^-/-^* mouse tissues.

Independent evidence also suggests that EFHD1 is primarily associated with the outer membrane protein. In a study using BioID proximity ligation to map mitochondrial proteomic interactions, EFHD1 was found as a bait primarily with BioID-tagged outer membrane proteins, such as *MAVS, MTCH2,* and *MFN2* [37]. Separately, a structural study revealed that EFHD1 alters cytoskeletal actin polymerization, which would accessible only from the outer membrane [9]. Finally, EFHD1 lacks a canonical mitochondrial targeting sequence, common among inner membrane and matrix proteins. Taken together, results here are best reconciled with prior evidence if EFHD1 primarily resides in the mitochondrial outer membrane and intermembrane space, with its reputed inner membrane localization possibly a consequence of looser associations from the intermembrane space with integral inner membrane proteins, rather than a restricted or constitutive inner membrane localization.

### EFHD1 ablation does not produce an overt cardiac phenotype, but is associated with reduced cardiac mitochondrial ROS levels and resistance to hypoxic death

We found no adverse effects of EFHD1 ablation on organismal growth or metabolism, cardiac physiology and function, and cardiac mitochondrial respiration. This stands in contrast to studies performed in sensory neurons and differentiating B cells, where a reduction in mitochondrial metabolism and shift towards glycolysis was noted [17, 18]. Part of the effect may be due to the levels of EFHD1, which are higher in neurons compared to cardiomyocytes, though even within the nervous system there were apparently minimal functional deficits.

The production of mitochondrial ROS is often a byproduct of respiratory flux and occurs mainly in the matrix and inner membrane. Yet, despite no change in cardiac mitochondrial respiration, we observed a robust reduction in mitochondrial ROS levels. Similarly, mitoflashes, the ROS-dependent respiratory bursts producing transient mitochondrial depolarization, were reduced by more than half in *Efhd1^-/-^* cardiomyocytes, with many cells producing none at all [7]. This result is consistent with prior data showing diminished mitoflash frequency in HeLa cells depleted of EFHD1 [6]. Furthermore, the discrepancy between reduced ROS despite unaltered respiration suggests that EFHD1 may be influencing the mitochondrial anti-oxidant response. Indirect evidence for this comes from prior studies showing that alterations in the levels of Superoxide Dismutase 2, an important mitochondrial enzyme for detoxifying excess superoxide anions, are countered by opposing changes in *EFHD1* expression [16]. The molecular mechanism by which EFHD1, residing outside the matrix, alters the function of mitochondrial ROS production, is likely indirect and remains to be determined, though the effect appears reproducible across several cell types.

Mitoflashes are thought to be precursors to the MPT, as excess ROS is a potent trigger for this event. We would therefore expect that mitochondria lacking *Efhd1* would be less susceptible to Ca^2+^-induced MPT. However, when performing the standard Ca^2+^ retention capacity assays, we found unchanged sensitivity in *Efhd1^-/-^* hearts (Figure 6A, B). This finding was surprising and suggests that mitochondrial sensitivity to ROS can be altered independently of its susceptibility to Ca^2+^ overload. One caveat is that CRC assays were performed on isolated mitochondria. If EFHD1 is involved with interactions between the mitochondrial outer membrane and other organelles, such as the ER, the limitations of studying the mitochondria in isolation needs to be taken into account.

Arguably the most profound phenotype which we report here is the resistance to CO_2_ euthanasia exhibited by *Efhd1*^-/-^ mice and the corresponding resistance to ischemia-reperfusion in isolated cardiomyocytes. This finding is intriguing, because it suggests the reduced ROS and mitoflash production in *Efhd1*^-/-^ cardiomyocytes may lead to a resistance to ischemic injury at the organismal level. Because our results have shown that EFHD1 ablation produces no obvious cardiac pathology, targeting this protein or its interactions may be a safe intervention providing a potentially novel therapy for cardiac injury.

### Limitations

Our intent in this study was to provide an initial characterization of the baseline cardiac phenotype in *Efhd1^-/-^* mice. It is unclear if the effects on ROS and cell death will translate to cardioprotection during different forms of cardiac injury. In addition, having determined that EFHD1 is primarily resident on the outer membrane and intermembrane space, the precise molecular mechanism by which it transduces the effects on ROS remains unexplained. Finally, given no change in susceptibility to mitochondrial Ca^2+^ overload, how the Ca^2+^-binding EF hands of this proteins alter its downstream effects will need to be established.

## DISCLOSURES

None

## SOURCES OF FUNDING

This work was supported by the Heart, Lung, and Blood Institute of the NIH under award R01HL141353 (D.C.), American Heart Association under Postdoctoral Fellowship award 834544(D.E.), Larry H. Miller Driving Out Diabetes Initiative (D.C.), and the Nora Eccles Treadwell Foundation (D.C.). The content is solely the responsibility of the authors and does not necessarily represent the official views of the funding agencies.

---

**Figure S1:**
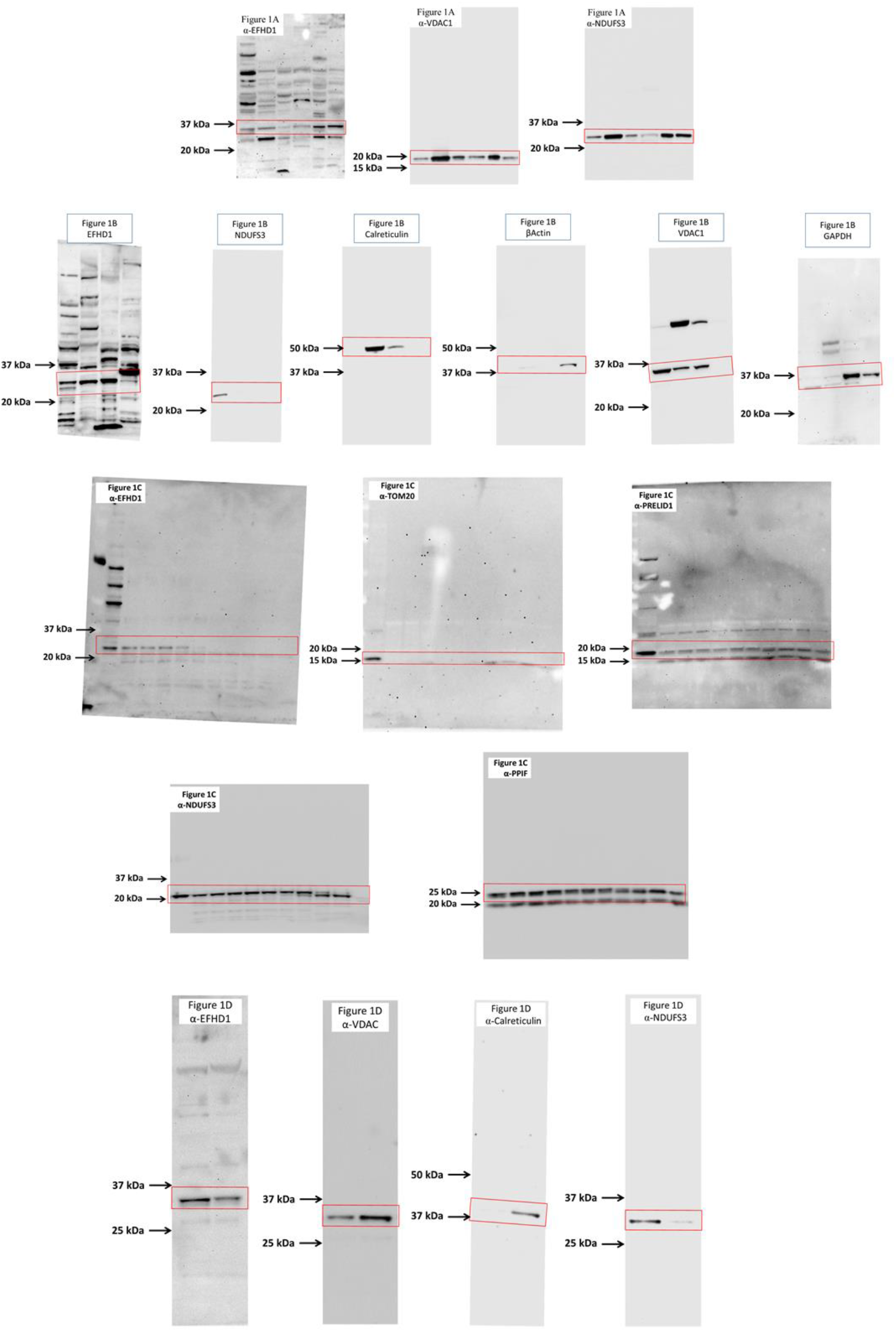
Expanded views of Western blots used in Figure 1.

**Figure S2:**
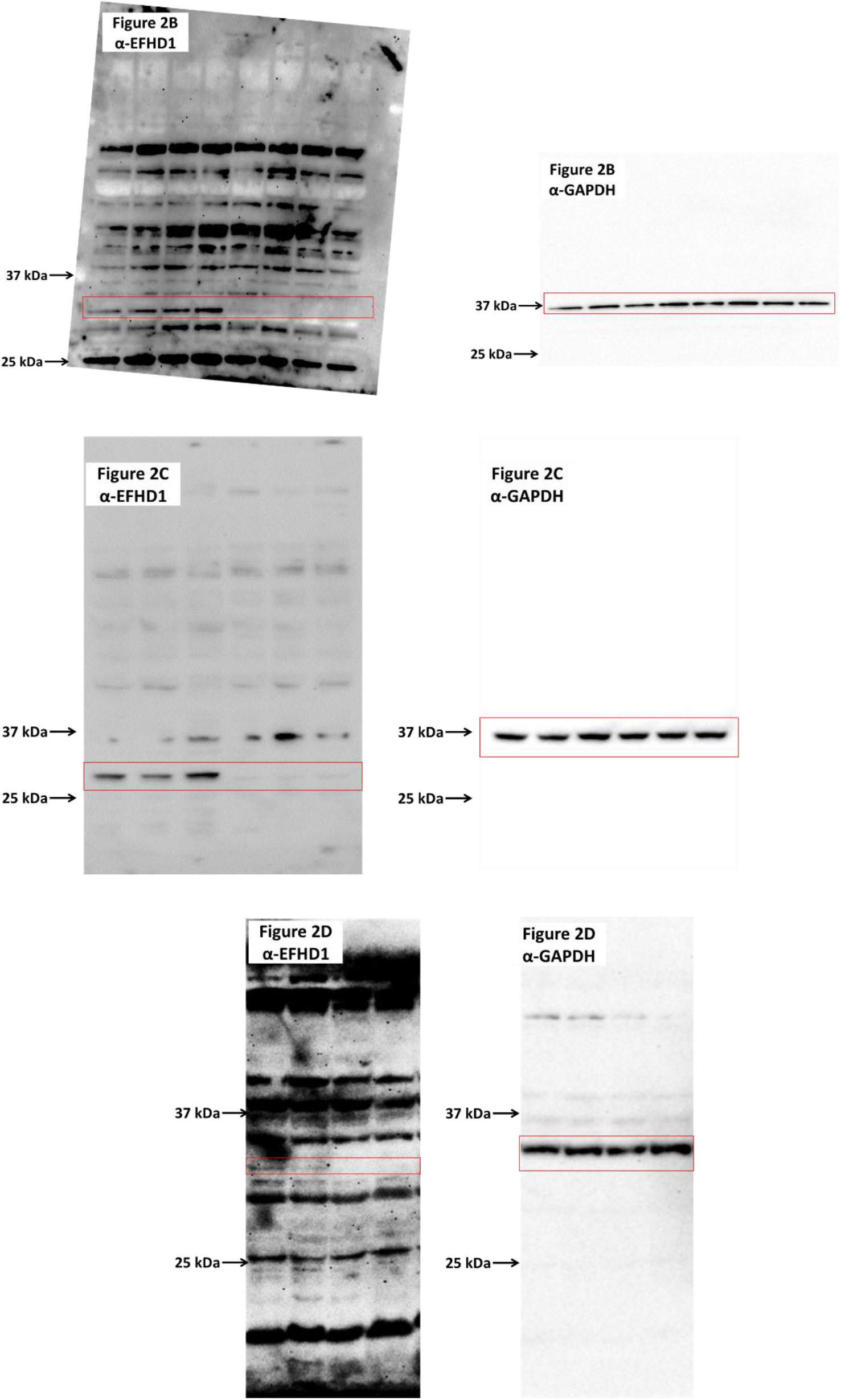
Expanded views of Western blots used in Figure 2.

**Figure S3:**
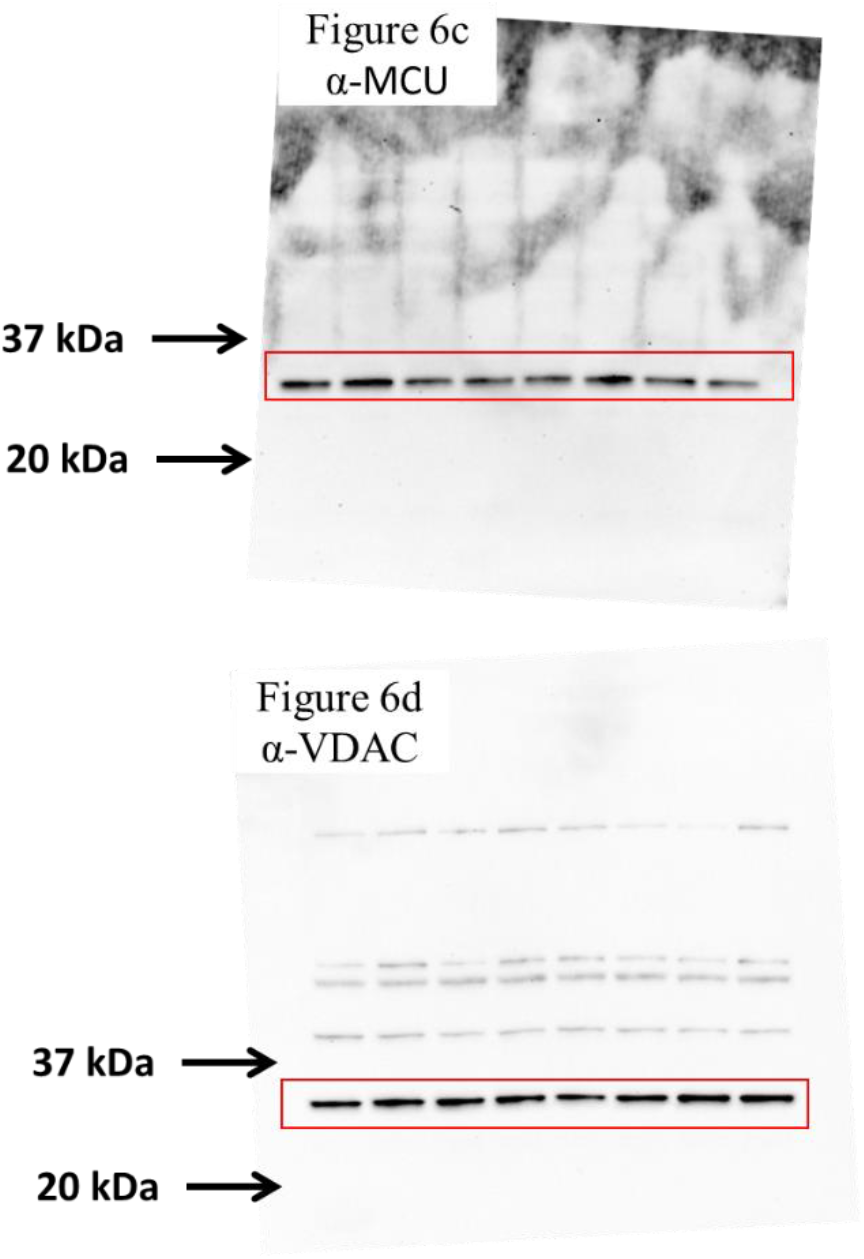
Expanded views of Western blots used in Figure 6.

## Notes

### Competing Interest Statement

The authors have declared no competing interest.

